# Rescuing AAV gene transfer from antibody neutralization with an IgG-degrading enzyme

**DOI:** 10.1101/2020.05.12.092122

**Authors:** Zachary C. Elmore, Daniel K. Oh, Katherine E. Simon, Marcos M. Fanous, Aravind Asokan

## Abstract

Pre-existing humoral immunity to recombinant adeno-associated viral (AAV) vectors restricts the treatable patient population and efficacy of human gene therapies. Approaches to clear neutralizing antibodies (NAbs), such as plasmapheresis and immunosuppression are either ineffective or cause undesirable side effects. Here, we describe a clinically relevant strategy to rapidly and transiently degrade NAbs prior to AAV administration using an IgG degrading enzyme (IdeZ). We demonstrate that recombinant IdeZ efficiently cleaves IgG in dog, monkey and human antisera. Prophylactically administered IdeZ cleaves circulating, human IgG in mice and prevents AAV neutralization *in vivo*. In macaques, a single intravenous dose of IdeZ rescues AAV transduction by transiently reversing seropositivity. Importantly, IdeZ efficiently cleaves NAbs and rescues AAV transduction in mice passively immunized with individual human donor sera representing a diverse population. Our antibody clearance approach presents a new paradigm for expanding the prospective patient cohort and improving efficacy of AAV gene therapy.

## Introduction

Human gene therapy using recombinant AAV vectors continues to advance steadily as a treatment paradigm for rare, monogenic disorders. This is highlighted by the recent FDA approval and clinical success of Zolgensma®, an intravenously dosed AAV vector delivering a functional copy of the *SMN1* gene in children with Spinal Muscular Atrophy (SMA)(1). Further, the list of systemically dosed AAV-based gene therapies for rare disorders such as Hemophilia A & B, Duchenne Muscular Dystrophy (DMD), X-linked myotubularin myopathy (XLMTM) and Pompe disease amongst others continues to grow(2, 3). These promising clinical examples have concurrently highlighted important challenges that include manufacturing needs, patient recruitment, and the potential for toxicity at high AAV doses. One such challenge that limits the recruitment of patients for gene therapy clinical trials and adversely affects the efficacy of AAV gene therapy is the prevalence of pre-existing neutralizing antibodies (NAbs) to AAV capsids in the human population. Such NAbs arise due to natural infection or cross-reactivity between different AAV serotypes(4–7). NAbs can mitigate AAV infection through multiple mechanisms by (a) binding to AAV capsids and blocking critical steps in transduction such as cell surface attachment and uptake, endosomal escape, productive trafficking to the nucleus or uncoating and (b) promoting AAV opsonization by phagocytic cells, thereby mediating their rapid clearance from the circulation. Multiple preclinical studies in different animal models have demonstrated that pre-existing NAbs impede systemic gene transfer by AAV vectors(8–11).

In humans, serological studies reveal a high prevalence of NAbs in the worldwide population, with about 67% of people having antibodies against AAV1, 72% against AAV2, and ∼ 40% against AAV serotypes 5 through 9(4, 12–14). Because of this high NAb sero-prevalence, screening for AAV antisera through *in vitro* NAb assays or ELISA is common place in AAV gene therapy trials and exclusion criteria can render upwards of 50% of patients ineligible for treatment or admission into clinical trials(15, 16). Furthermore, vector immunogenicity represents a major challenge in re-administration of AAV vectors. High titer NAbs are produced following AAV vector administration, thereby preventing prospective AAV redosing(6, 17). This severely limits long term gene therapy success in (a) patients in the low dose AAV cohort; (b) pediatric patients who will experience tissue growth and proliferation leading to vector genome dilution and potential reversal of symptoms with age, and (c) patients with degenerative disorders that might require multiple AAV treatments to prevent tissue loss and sub-therapeutic transgene expression levels. Taken together, NAbs present a significant barrier to the broad application of AAV in the clinic.

Strategies that are currently being evaluated to circumvent pre-existing humoral immunity to AAV vectors are either early in development, ineffective or prone to causing undesirable side effects. These include the engineering of new AAV variants with reduced NAb recognition(18, 19), plasmapheresis or immunoadsorption to reduce the overall levels of circulating antibodies in patient serum prior to AAV administration(20–23), use of capsid decoys(24) or immunosuppression to decrease the B cell population and consequently antibody levels in general(25, 26). While these approaches have demonstrated varying success and efficiency in addressing the problem of circulating antibodies and remain under evaluation, a one-solution-fits-all approach that resolves this challenge is unlikely. Pertinent to this, a promising and clinically validated paradigm for mitigating the effects of deleterious (auto)antibodies is the use of IgG-specific proteases(27–30). In particular, the extracellular enzyme, IdeS derived from *Streptococcus pyogenes*, is a 35 kDa cysteine protease that specifically cleaves IgG at the lower hinge region generating one F(ab′)_2_ fragment and one homodimeric Fc fragment(31–34) (**Figure 1A**). IdeZ, a homolog of IdeS, was identified and characterized in *S. equi* ssp. *zooepidemicus* and shown to efficiently cleave IgG in a similar manner to IdeS(35, 36). Here, we evaluate the ability of IdeZ to mitigate the effect of pre-existing anti-AAV NAbs in mice passively immunized with human antisera and in non-human primates. First, we demonstrate the ability of IdeZ to cleave antibodies in sera derived from multiple species. Next, we show that IdeZ can rescue AAV gene transfer in the presence of circulating human IgG in mice and natural humoral immunity in non-human primates. In addition, we demonstrate that gene transfer to the liver and heart is also rescued in mice passively immunized with individual human antisera.

**Figure 1.**
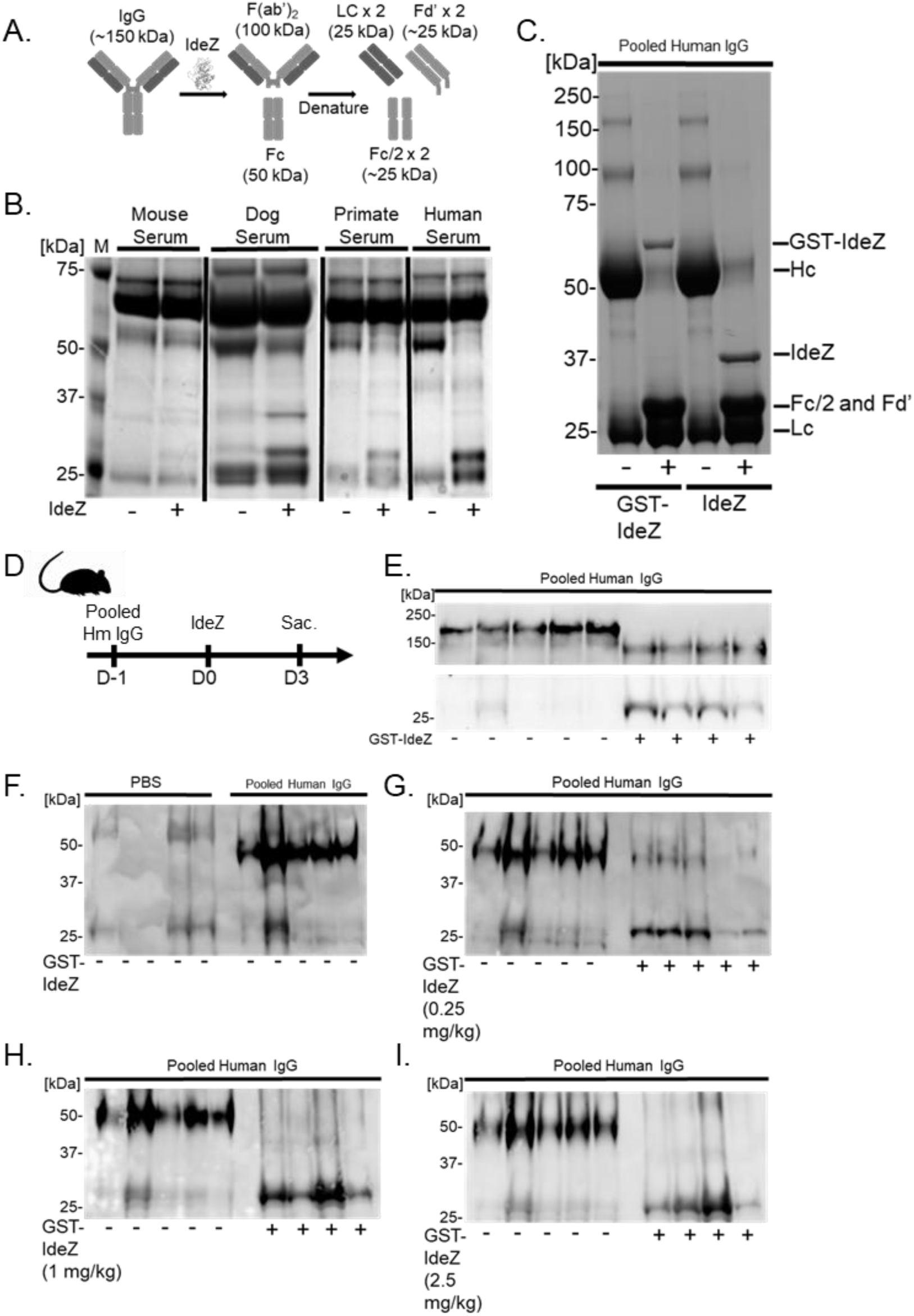
IdeZ cleaves serum antibodies from multiple species. **A**, Schematic outlining IdeZ cleavage of IgG below the hinge region yielding multiple F(ab’)_2_ and Fc fragments after reduction. **B**, Serum samples from mouse, dog, primate and human untreated (-) or treated (+) with recombinant IdeZ and analyzed by SDS-PAGE under reducing conditions. Gels were then stained with Coomassie blue. **C**, Pooled human IgG untreated (-) or treated (+) with recombinant GST-IdeZ or commercial standard IdeZ (NEB) and analyzed by SDS-PAGE under reducing conditions. Gels were then stained with Coomassie blue. IgG was cleaved by GST-IdeZ and IdeZ into multiple fragments as indicated. **D**, Mice were injected intraperitoneally first with pooled human IgG, following which they were injected intravenously 24 hours later with PBS (-) or recombinant GST-IdeZ (1 mg/kg) (+). Blood samples were taken 72 hours post injection and analyzed by SDS-PAGE under reducing conditions with immunoblotting. IgG was probed with Fab (top panel) and Fc (bottom panel) specific antibodies. **E**, Experimental timeline of *in vivo* GST-IdeZ dose optimization experiment. Mice were injected with pooled human IgG followed 24 hrs later with no injection or injection with 3 different doses of GST-IdeZ. Blood serum samples were collected 72 hours post GST-IdeZ. Sac., sacrifice followed by tissue harvest. Serum samples of PBS control (**F**), 0.25 mg/kg (**G**), 1 mg/kg (**H**), and 2.5 mg/kg (**I**) GST-IdeZ injected mice were analyzed by SDS-PAGE under reducing conditions and probed with human IgG specific antibodies to analyze IgG cleavage.

## Results

### IdeZ shows robust ability to cleave antibodies in sera from multiple species

We first demonstrated that IdeZ efficiently cleaves antibodies in canine, non-human primate and human sera, but not mouse serum samples *in vitro* (**Figure 1B**). The latter observation is corroborated by known mutations in the hinge region of mouse IgG compared to other species that render the latter resistant to IdeZ mediated degradation(35, 36). IdeZ also effectively cleaves human IgG into heavy chain, light chain and Fc fragments *in vitro* (**Figure 1C**). Next, we confirmed the potency of research grade, recombinant GST-tagged IdeZ produced in *E.coli* for dosing *in vivo* (**Figure 1D**). Mice were first passively immunized with pooled IgG injected intraperitoneally (IP), followed by a single intravenous (IV) injection of IdeZ confirming efficient cleavage into Fab and Fc fragments as determined by western blotting (**Figure 1E)**. Further, as shown in **Figures 1F-I**, we observed a dose dependent effect in IgG degradation, with optimal clearance between 0.25-1mg/kg of IdeZ at day 2-3 post-administration. Effective clearance of circulating antibodies was observed within a day or two post-IdeZ administration **(Supplemental Figure S1).** Furthermore, IdeZ effectively mitigated human IgG mediated neutralization of AAV8 and AAV9-Luc transduction *in vitro* (**Supplemental Figure S2**) leading us to investigate the efficacy of IdeZ treatment on AAV gene transfer efficiency in the presence of neutralizing antisera *in vivo*.

### IdeZ rescues AAV liver gene transfer in mice and macaques

Based on these results, we evaluated the ability of prophylactically dosing IdeZ in mice passively immunized with pooled human IgG to rescue AAV transduction *in vivo*. Briefly, animals of either gender were first injected IP with pooled human IgG (8 mg) on day (−1), with a single dose of IdeZ (2.5mg/kg) through the tail vein on day 0 and an IV dose of AAV8 or AAV9 vectors (1×10^13^vg/kg) packaging a CBA promoter driven luciferase transgene on day 3 (**Figure 2A**). Naïve mice showed different levels of AAV8 and AAV9-mediated luciferase expression in the liver (**Figures 2B-E**). In mice passively immunized with pooled human IgG, luciferase expression in the liver was decreased by 10-100 fold due to the presence of anti-AAV NAbs. In contrast, we observed rescue from AAV neutralization in IdeZ treated animals, with partial to complete rescue of liver luciferase expression levels. These observations were corroborated by vector genome copy numbers, which corresponded with transgene expression in general; although, we observed gender-specific differences (Figures 2F-I). Notably, despite restoration of AAV copy numbers in the male liver, expression was not fully restored implying that other non-NAb related factors might be involved in controlling liver expression (**Figure 2C,E**). While these aspects warrant further investigation and dose optimization, these observations support that prophylactically administered IdeZ can prevent AAV neutralization and restore liver transduction in an AAV serotype-independent manner.

**Figure 2.**
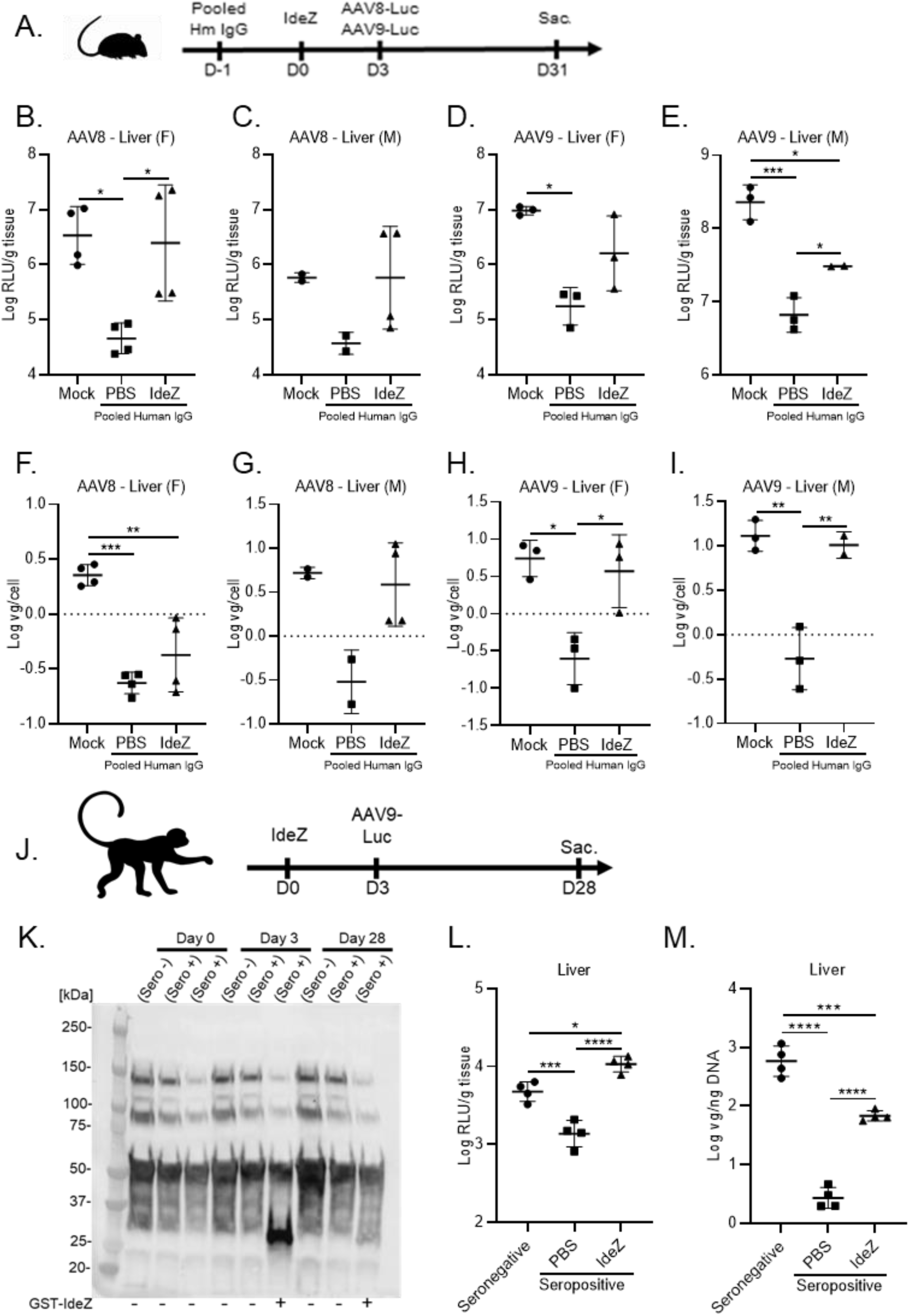
IdeZ rescues AAV8 and AAV9 liver transduction in passively immunized mice and cynomolgus macaques. **A**, Experimental timeline of IgG, IdeZ and AAV8 or AAV9-Luc injections. Sac., sacrifice followed by tissue harvest. Mice were injected intraperitoneally with pooled human IgG. The same mice were injected intravenously 24 hours later with PBS or recombinant GST-IdeZ (2.5 mg/kg). AAV8-Luc or AAV9-Luc was injected 72 hours post IdeZ at a dose of 1 × 10^13^ vg/kg. Luciferase transgene expression levels were analyzed 4 weeks post-injection in the liver; AAV8 (**B,C**); AAV9 (**D,E**). Luciferase expression levels were normalized for total tissue protein concentration and represented as log relative luminescence units per gram of tissue (log RLU/g tissue). Each dot represents the average of a technical duplicate from a single animal. Biodistribution of AAV8 and AAV9 Luc vector genomes in the liver; AAV8 (**F,G**); AAV9 (**H,I**). Vector genome copy numbers per cell were calculated by normalizing Luc copy numbers to copies of the Lamin B2 housekeeping gene and represented as log vg/cell. Each dot represents a technical duplicate from a single animal, and the dash represents the mean value. (F=female, M=male). **J**, Schematic demonstrating experimental timeline of IdeZ and AAV9-Luc injections in NHPs. AAV9 seropositive NHP M16558 (n=1) was administered IdeZ (0.5 mg/kg) via intravenous bolus injection on Day 0. AAV9-Luc was administered via intravenous bolus injection 72 hrs post-IdeZ injection at a dose of 5 × 10^12^ vg/kg. **K**, NHP serum samples were analyzed by SDS-PAGE under reducing conditions and probed with Fc specific antibodies. **L**, Luciferase transgene expression levels were analyzed 4 weeks post-injection in the liver of NHPs. Luciferase expression levels were normalized for total tissue protein concentration and represented as log relative luminescence units per gram of tissue (Log RLU/g tissue). Each dot represents a single experiment of an individual liver lobe from a single animal. **M**, Biodistribution of AAV9 Luc vector genomes in the liver of NHPs. Vector genome copy numbers per ng of total extracted DNA were calculated and represented as log vg/ng DNA. Each dot represents a technical duplicate experiment of individual liver slices from a single animal and the dash represents the mean value. Significance was determined by one-way ANOVA with Tukey’s post-test. \**p*<0.05, \*\**p*<0.01, \*\*\**p*<0.001, \*\*\*\**p*<0.0001

Next, we sought to evaluate whether IdeZ was effective in non-human primates. We first screened male cynomolgus macaques for anti-AAV antibodies using a NAb assay to identify seropositive and seronegative animals (**Supplemental Figure S3**). Animal M16561 served as the naïve seronegative control, while the seropositive animals M16556 and M16558 were dosed on day 0 with IV PBS or a single IV bolus dose of IdeZ (0.5mg/kg), respectively. On day 3, post-IdeZ treatment, all three animals were injected with a dose of AAV9 vectors packaging the luciferase transgene (5×10^12^ vg/kg) (**Figure 2J**). Evaluation of serum IgG levels at days 0, 3 and 31 post-IdeZ treatment revealed selective cleavage and clearance at day 3. In addition, serum IgG levels were fully restored to normal levels by day 31 corroborating the transient effect of IdeZ activity (**Figure 2K**). Upon sacrifice at day 30, we observed an approximately one log order decrease in luciferase gene expression and a disproportionate (∼ two logs) decrease in vg copy number in the liver (**Figure 2L & 2M**). Importantly, IdeZ treatment restored AAV luciferase gene expression levels and partially restored vg copy numbers in the liver. Further, these results also mirrored the observations in the liver of male mice injected with human IgG. While it should be noted that the number of non-human primates in the current study are low, the above results underscore the ability to translate the applicability of IdeZ in clearing IgG across multiple species.

### IdeZ rescues AAV liver gene transfer in mice passively immunized with individual human sera

To further evaluate whether IdeZ can function effectively in a clinically relevant setting, we tested our approach in mice passively immunized with individual human donor sera representing a diverse population. Briefly, we obtained 18 different human donor serum samples across a broad demographic and displaying varying levels of AAV neutralization as determined by NAb assay (**Supplemental Figure S4**). We then administered a single IP dose of donor serum in 2 animals each (total 18 cohorts), following which the first animal received an IV injection of PBS and the second, a single IV bolus dose of IdeZ (0.5mg/kg). The control cohort comprised of naïve mice. All animals received an IV dose of AAV9-Luc vectors (1×10^13^ vg/kg) and luciferase gene expression assessed in the liver and heart of the saline vs IdeZ treated cohorts (**Figure 3A**). As seen in **Figures 3B-E**, the diversity of pre-existing humoral immunity to AAV transduction is well represented by this small, yet diverse panel of human serum samples (**Supplemental Table 1**). Notably, we observed restoration in liver luciferase expression levels in a number of animals (**Figures 3B,D**). Complete restoration (100%) of liver expression to that of naïve, non-immunized control animals was observed in these animals regardless of NAb titer. Some outliers were also observed, where IdeZ treatment was only partially effective or adversely affected transduction. One possible explanation is that these mice might have high levels of pre-existing immunity to IdeZ, although the impact of such on IdeZ activity is unclear. While these aspects warrant further investigation, we observed overall trends that support that IdeZ treatment can result in a statistically significant improvement in liver gene expression and copy number by clearing circulating antibodies (**Figures 3C,E**). These results further underscore the potential for clinical translation with our approach.

**Figure 3.**
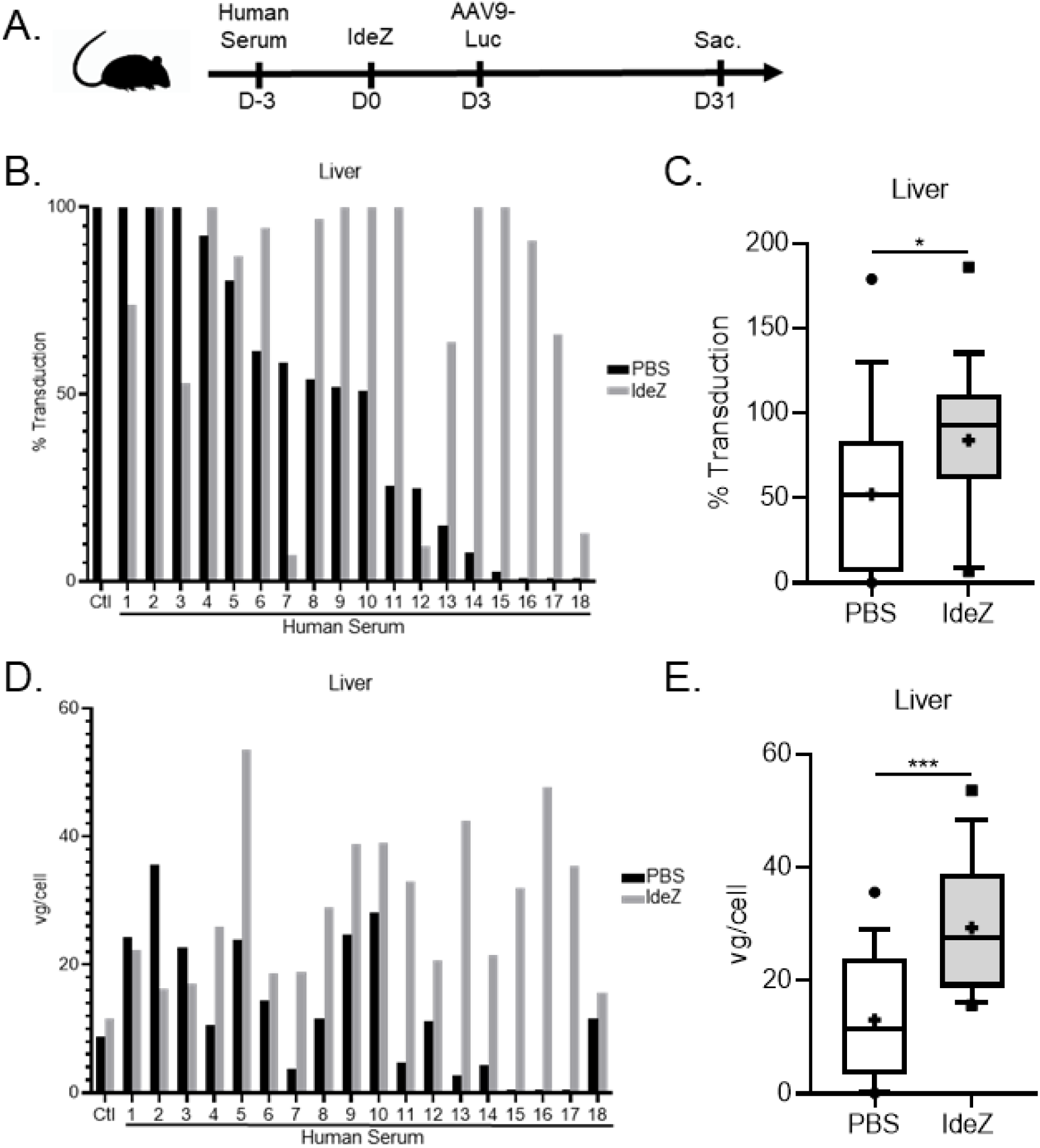
IdeZ rescues AAV9 liver transduction in mice passively immunized with individual human sera. **A**, Schematic demonstrating experimental timeline of human serum, IdeZ and AAV9-Luc injections. 18 human serum samples were tested for their ability to neutralize AAV9 transduction in the liver. Two mice per human serum sample were utilized for the study and both mice were injected intraperitoneally with human serum. Mice were then injected intravenously 72 hours later with PBS (black bars) or recombinant GST-IdeZ (2.5 mg/kg, grey bars) and subsequently injected intravenously 72 hrs post-IdeZ treatment with AAV9-Luc (1 × 10^13^ vg/kg). Liver transduction levels were analyzed 4 weeks post-injection. Sac., sacrifice followed by tissue harvest. **B**, Luciferase transgene expression levels were analyzed 4 weeks post-injection in the liver of passively immunized mice treated with PBS (black) or prophylactically with IdeZ (grey). Transduction levels were normalized to non-immunized mice that were injected with AAV9-Luc at the same dose and represented as percentage of control. Each bar represents the average of a technical duplicate from a single animal. **C**, Relative liver transduction efficiency of AAV9-Luc in the entire cohort of mice immunized with human sera treated with PBS control (white) or IdeZ (grey). Biodistribution of AAV9 vector genomes in the liver for mice passively immunized with individual human serum samples (**D**) and the entire cohort (**E**). Vector genome copy numbers per cell were calculated based on normalization to copies of the Lamin B2 housekeeping gene. Each bar represents the average of a technical duplicate from a single animal. Significance was determined by the nonparametric Mann-Whitney rank test. \**p*<0.05, \*\**p*<0.01, \*\*\**p*<0.001, \*\*\*\**p*<0.0001.

### IdeZ mediated rescue of AAV cardiac gene transfer efficiency provides additional insight into plausible neutralization mechanisms

Concurrent to studies focused on restoring AAV gene transfer in the liver, we also analyzed the heart and observed striking differences. We assessed cardiac gene transfer in both mice passively immunized with pooled human IgG as well as individual human sera. Notably, although pooled human IgG decreased expression and IdeZ treatment restored cardiac luciferase expression levels to that of naïve mice, changes in vg copy number upon IdeZ treatment were only partially rescued in females or statistically insignificant in males (**Figures 4A-E**). These observations were further corroborated in mice passively immunized with individual human sera. In this regard, we first observed that neutralization of cardiac transduction by individual antisera does not mirror the patterns observed in the liver (**Figures 3B,D** and **4G,I**, black columns). Second, only partial rescue of AAV mediated cardiac gene expression is observed in most animals. In addition, although we observed some increase in vector genome copy numbers within cardiac tissue, no specific correlation with luciferase expression patterns was noted (**Figures 4G,I**). Assessment of overall rescue across the human sera infused cohorts corroborated these trends (**Figures 4H,J**). In particular, we observed a statistically significant rescue of cardiac gene transfer in cardiac luciferase expression, but not vector genome copy numbers.

**Figure 4.**
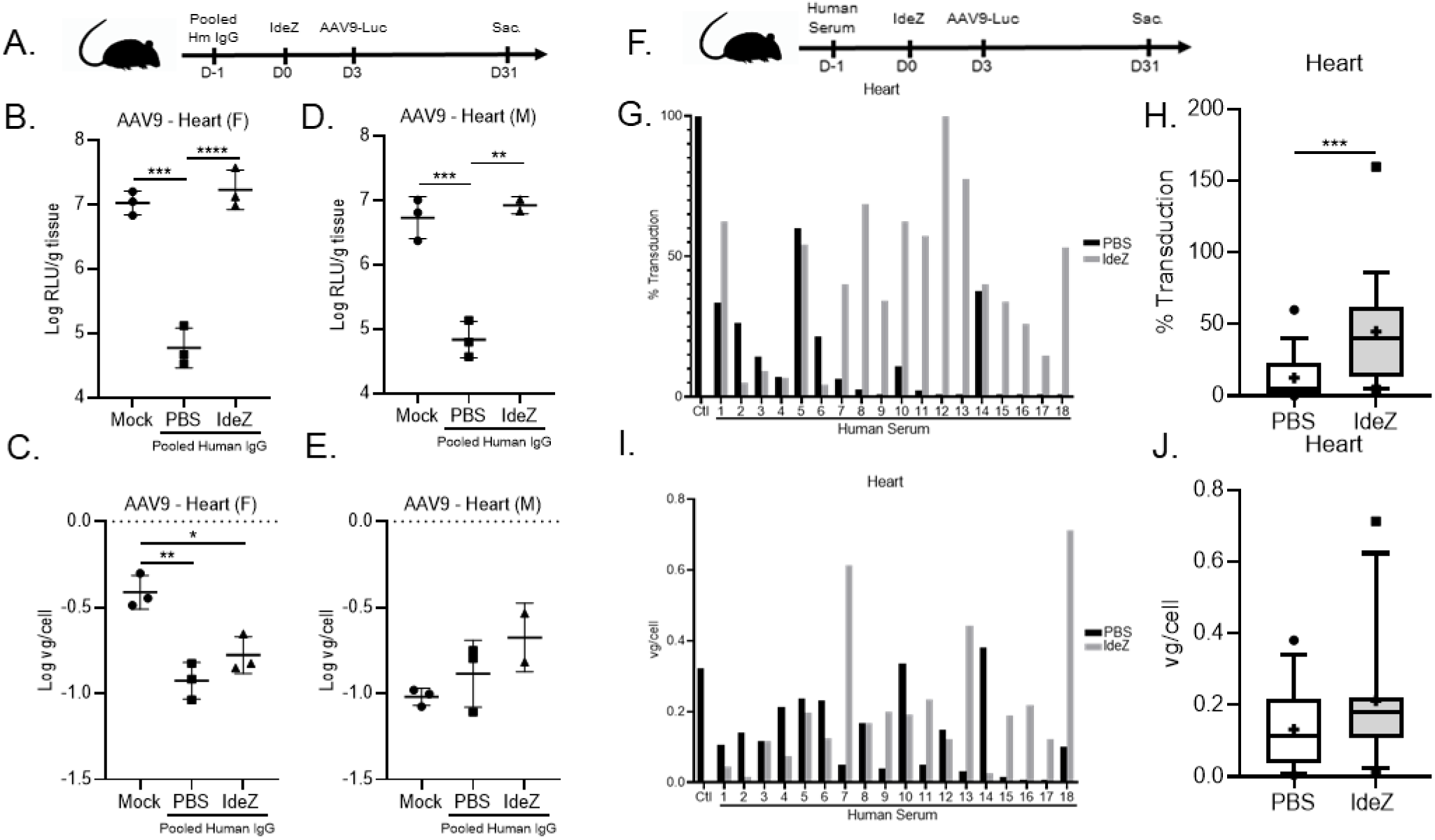
Impact of IdeZ treatment on AAV9 cardiac transduction in passively immunized mice. **A**, Experimental timeline of pooled human IgG, IdeZ and AAV9-Luc injections. Cardiac tissues were derived as outlined earlier in the liver experiment. **B,D**, Luciferase transgene expression levels were analyzed 4 weeks post-injection in the heart. Luciferase expression levels were normalized for total tissue protein concentration and represented as log relative luminescence units per gram of tissue (log RLU/g tissue). Each dot represents the average of a technical duplicate from a single animal. **C,E**, Biodistribution of AAV9 Luc vector genomes in the heart. Vector genome copy numbers per cell were calculated based on normalization to copies of the Lamin B2 housekeeping gene. Each dot represents the average of a technical duplicate from a single animal. Female (F), Male (M). Significance was determined one-way ANOVA with Tukey’s post-test. \**p*<0.05, \*\**p*<0.01, \*\*\**p*<0.001, \*\*\*\**p*<0.0001. **F**, Schematic demonstrating experimental timeline of human serum, IdeZ and AAV9-Luc injections. Cardiac tissues were derived as outlined earlier in the liver experiment. **G**, Luciferase transgene expression levels were analyzed 4 weeks post-injection in the heart of passively immunized mice treated with PBS (black) or prophylactically with IdeZ (grey). Transduction levels were normalized to non-immunized mice that were injected with AAV9-Luc at the same dose and represented as percentage of control. Each bar represents the average of a technical duplicate from a single animal. **H**, Relative cardiac transduction efficiency of AAV9-Luc in the entire cohort of mice immunized with human sera treated with PBS control (white) or IdeZ (grey). Biodistribution of AAV9 vector genomes in the heart for mice passively immunized with individual human serum samples (**I**) and the entire cohort (**J**). Vector genome copy numbers per cell were calculated based on normalization to copies of the Lamin B2 housekeeping gene. Each bar represents the average of a technical duplicate from a single animal. Significance was determined by the nonparametric Mann-Whitney rank test. \**p*<0.05, \*\**p*<0.01, \*\*\**p*<0.001, \*\*\*\**p*<0.0001.

## Discussion

The IgG-degrading enzyme, IdeS, also known as Imlifidase® has shown promise in a clinical trial (ClinicalTrials.gov Identifier: NCT02224820) permitting successful kidney transplantation in patients harboring donor-specific antibodies(37–39). Briefly, the latter study assessed the safety, immunogenicity, pharmacokinetics, and efficacy of Imlifidase in an open-label, dose escalation study in highly sensitized patients with anti-HLA antibodies and chronic kidney disease. This approach represents a potential paradigm shifting method to desensitize patients, who would otherwise not qualify to receive a lifesaving transplant. Thus, a clinical precedent for applying enzymatic IgG degradation to promote rapid and transient antibody clearance already exists. Further, it is noteworthy to mention that other orthogonal methods to facilitate IgG clearance using soluble antibody binding bacterial proteins (e.g., Protein M(40)), FcRn (neonatal Fc receptor) domains(41), anti-FcRn antibodies such as Rozanolixizumab(42), SYNT001(43, 44) etc have shown promise in the clinic as well.

These approaches, however, have not been explored in the context of gene therapy to date. The antibody degradation/clearance approach described in the current study could broadly impact preclinical gene therapy studies in different large animal models, currently encumbered by pre-existing NAbs. For instance, pre-existing humoral immunity against different AAV serotypes in macaques, dogs and pigs have been described(8, 10). Based on our earlier *in vitro* results, we postulate that IdeZ could potentially be applicable for evaluating AAV gene therapies in canine models of disease. These data combined with our observations in non-human primates greatly expands the potential for preclinical AAV gene transfer studies, but also provides a path towards safety and dose finding studies of this approach in preclinical animal models. Additional studies to evaluate IdeZ dosing and kinetics of antibody clearance in such animal subjects with varying anti-AAV antibody titers is likely to help optimize this approach.

Another important advantage of the IdeZ approach is the potential for AAV serotype-independent rescue from antibody neutralization. While such will require dose optimization studies with different natural and engineered AAV capsids, we postulate that the universality of our antibody clearance approach is likely to broadly complement AAV gene transfer studies. One possible caveat of this approach is that people may harbor antibodies against IdeZ. However, it is interesting to note that IdeZ would likely degrade such antibodies as well. Another significant topic that warrants further evaluation is whether IdeZ treatment can enable vector redosing. In particular, IdeZ could provide an alternative solution in patients, where immunosuppression is not feasible or undesirable(25, 26, 45). While we were unable to evaluate such in mice due to the inability of IdeZ to cleave mouse IgG, such studies should be feasible in non-human primates or other animal models. Taken together, from a clinical perspective, the current strategy has the potential to significantly impact the treatable patient population and improve the efficacy of AAV gene therapies.

## Methods

### Plasmid Constructs and Recombinant Protein Expression

IdeZ DNA sequence from *S. equi* ssp. *zooepidemicus* lacking the N-terminal signal seqence was synthesized and cloned into pGEX-6P-3 expression vector using BamHI and SalI restriction sites (Genscript). *E. coli* strain BL21 star (DE3) was transformed with recombinant IdeZ pGEX-6P-3 plasmid. A single colony was inoculated into TB medium containing ampicillin; culture was incubated in 37°C at 200 rpm and then induced with IPTG. Recombinant BL21 cells stored in glycerol were inoculated into TB medium containing ampicillin and cultured at 37 °C. When the OD600 reached about 4, the cell culture was induced with IPTG at 37°C for 4h. Cells were harvested by centrifugation. Cell pellets were resuspended with GST lysis buffer followed by sonication. The supernatant after centrifugation was kept for future purification. Target protein was obtained by two-step purification using a GST column and Superdex 200 column. Target protein was sterilized by 0.22μm filter before stored in aliquots. The concentration was determined by Bradford protein assay with BSA as standard. The protein purity and molecular weight were determined by standard SDS-PAGE along with Western blot confirmation using a Rabbit anti-GST pAb (GenScript, Cat.No. A00097). Recombinant GST-IdeZ was stored in 50 mM Tris-HCl, 150 mM NaCl, 10% Glycerol, pH 8.0. Endotoxin was removed from recombinant protein using High Capacity Endotoxin Removal Spin Columns (ThermoFisher Scientific Catalog #88274) following the manufacturer’s instructions.

### SDS-PAGE and analysis of IdeZ enzyme activity

Pooled human IgG was purchased from Sigma (I4506), Mouse and Dog serum samples were obtained from lab stocks or kind gifts from David Mack (University of Washington). Individual human serum samples from donors were purchased from ValleyBiomedical. Rhesus macaque sera were kind gifts from Alice Tarantal (UC Davis), Yoland Smith and Adriana Galvan (Yerkes National Primate Center, Emory University). Proteins analyzed by SDS-PAGE were separated under reducing conditions on Nu-PAGE 4–12% Bis-Tris (Invitrogen) or on Mini-Protean TGX 4-15% gels (Biorad) and stained with Coomassie Blue. All *in vitro* activity assays with recombinant GST-IdeZ or IdeZ (NEB Catalog #P0770S) (1ug per reaction) were performed for 3 h at 37°C and serum samples were diluted 50x in PBS prior to analysis by SDS-PAGE. All *in vivo* activity assays were performed with recombinant GST-IdeZ with mouse and or NHP serum samples being diluted 10x in PBS prior to analysis by SDS-PAGE and immunoblotting. Digested sera was probed with Rabbit anti-Human IgG-HRP H+L Secondary Antibody (ThermoFisher Scientific Catalog #A18903, 1:10,000), Rabbit anti-Human IgG Fc HRP Secondary Antibody (ThermoFisher Scientific Catalog #31423, 1:10,000), and Rabbit anti-Human IgG F(ab’)_2_ HRP Secondary Antibody (ThermoFisher Scientific Catalog #31482, 1:10,000).

### Cell lines and Recombinant Virus Production

HEK293 were maintained in Dulbecco’s Modified Eagle’s Medium (DMEM) supplemented with 10% fetal bovine serum (FBS), 100U/ml penicillin, 100ug/ml streptomycin. Cells were maintained in 5% CO_2_ at 37°C. Recombinant AAV vectors were generated using triple plasmid transfection with the AAV Rep-Cap plasmid (pXR8 or pXR9 encoding AAV8 or AAV9 capsid proteins, respectively), Adenoviral helper plasmid (pXX680), and a luciferase transgene cassette driven by the chicken beta actin promoter (pTR-CBA-Luc), flanked by AAV2 inverted terminal repeat (ITR) sequences. Viral vectors were harvested from media and purified via iodixanol density gradientµltracentrifugation followed by phosphate-buffered saline (PBS) buffer exchange. Titers of purified virus preparations were determined by quantitative PCR using a Roche Lightcycler 480 (Roche Applied Sciences, Pleasanton, CA) with primers amplifying the AAV2 ITR regions (forward, 5′-AACATGCTACGCAGAGAGGGAGTGG-3′; reverse, 5′-CATGAGACAAGGAACCCCTAGTGATGGAG-3′) (IDT Technologies, Ames IA).

### *In Vitro* Antibody and Serum Neutralization Assays

Pooled human IgG, (25 ug undiluted) or antiserum (25µl) (as specified for individual experiments) was mixed with an equal volume containing recombinant AAV9-Luc vector (100,000 vg/cell) in tissue culture-treated, black, glass-bottom 96-well plates (Corning) and then incubated at room temperature for 30 min. For neutralization assays, 1 ug of GST-IdeZ was incubated with pooled human IgG in 5% CO2 at 37 °C for 2 h prior to addition of AAV9-CBA-Luc vector. A total of 1×10^4^ HEK293 cells in 50μL DMEM + 10% FBS + penicillin-streptomycin was then added to each well, and the plates were incubated in 5% CO2 at 37 °C for 24h. Cells were then lysed with 25μL of 1×passive lysis buffer (Promega) for 30 min at room temperature. Luciferase activity was measured on a Victor 3 multi-label plate reader (PerkinElmer) immediately after the addition of 25μL of luciferin (Promega). All readouts were normalized to controls with no antibody/antiserum treatment. Recombinant AAV vectors packaging CBA-Luc transgenes, antibodies, sera, and GST-IdeZ were prediluted in DMEM and used in this assay.

### Mouse Studies

All animal experiments were performed using 6- to 8-week-old male and female C57BL/6 mice purchased from Jackson Laboratories (Bar Harbor, ME). These mice were maintained and treated in compliance with NIH guidelines and as approved by the UNC Institutional Animal Care and Use Committee (IACUC). Mice were injected intraperitoneally with pooled human IgG (8 mg). The same mice were injected intravenously 24 hours later with PBS or recombinant GST-IdeZ (2.5 mg/kg). Recombinant AAV9-CBA-Luc or 1× PBS (as mock treatment) was injected 72 hours post IdeZ at a dose of 1 × 10^13^ vg/kg. Luciferase transgene expression levels were analyzed 4 weeks postinjection in the liver, heart, and kidney. Animals were sacrificed 4 weeks post-AAV9 injection with an intraperitoneal injection of tribromoethanol (Avertin) (0.2 ml of 1.25% solution) followed by transcardial perfusion with 30 ml of 1× PBS. For human serum samples/IdeZ studies, two mice were injected intraperitoneally with 200µl human sera (purchased from Valley Biomedical, gift from StrideBio, Inc.). The same mice were then injected intravenously 72 hours later with PBS or recombinant GST-IdeZ (2.5 mg/kg). Mice were subsequently injected intravenously 72 hrs post-IdeZ treatment with AAV9-Luc (1 × 10^13^ vg/kg).

### Non-human primate studies

A total of 3 cynomolgus macaques (3 males) designated for use in this study were obtained from Southern Research (Birmingham, AL) who obtained them from Worldwide Primates, Inc. (Miami, FL). Animals were acclimated prior to study start and deemed healthy prior to study initiation. On the first day of dosing, the animals were approximately 3 years of age, male gender and weighed between 2.4 – 3.8 kg. Housing and animal care conformed to the guidelines of the U.S. Department of Agriculture (Animal Welfare Act; Public Law 99-198) and those of the *Guide for the Care and Use of Laboratory Animals* and to the applicable Standard Operating Procedures (SOPs) at Southern Research. Animals were tested for pre-existing AAV9 NAbs using an *in vitro* NAb assay and were designated as M16561 being seronegative, M16556 being seropositive and M16558 being seropositive. The seropositive NHP M16558 was administered IdeZ (0.5 mg/kg) via intravenous bolus injection on Day 0. AAV9-CBA-Luc was administered to all 3 NHPs via intravenous bolus injection, 72 hrs post-IdeZ injection at a dose of 5 × 10^12^ vg/kg. All animals had blood collected for analysis on days 0, 3, and 28. On day 28, NHPs were euthanized and organs collected via whole body perfusion with sterile saline while under anesthesia following collection of specified blood samples.

### Tissue analysis for luciferase expression

To quantify luciferase expression, animals injected with AAV9-CBA-Luc transgene were sacrificed and tissues were harvested and frozen at 80C. Tissues were later thawed, weighed, and lysed by adding 200µl of 1x passive lysis buffer (Promega, Madison WI) prior to mechanical lysis using a Tissue Lyser II 352 instrument (QIAGEN, Valencia, CA), followed by centrifugation to remove any remaining tissue debris. To measure luciferase transgene expression, 15µl of supernatant from each lysate was then loaded onto an assay plate along with 45µl of luciferin, and luminometric analysis was performed using a Victor3 luminometer (PerkinElmer, Waltham, MA). The relative luminescence units obtained for each sample were then normalized to the input tissue weight for each sample, measured in grams, followed by log transformation.

### Tissue analysis for vector genome biodistribution

DNA was extracted and purified from tissues using a QIAamp DNA FFPE Tissue kit (catalog no. 56404; Qiagen). Viral genome copy numbers were then determined for each tissue using quantitative PCR with primers specific to the chicken beta actin (CBA) promoter (forward, 5′-TGTTCCCATAGTAACGCCAA-3′; reverse, 5′-TGCCAAGTAGGAAAGTCCCAT-3′). These viral genome copy numbers were then normalized to the level of the mouse lamin B2 housekeeping gene using specific primers (forward, 5′-GGACCCAAGGACTACCTCAAGGG-3′; reverse, 5′-AGGGCACCTCCATCTCGGAAAC-3′). The biodistribution of viral genomes is represented as the ratio of vector genomes per cell or as vector genomes per nanograms of DNA extracted, followed by log transformation.

### Statistical Analysis

Where appropriate, data are represented as mean or mean ± standard deviation. Where appropriate data were log transformed prior to statistical analysis. For data sets with at least three groups, significance was determined by one-way ANOVA, with Tukey’s post-test. For analysis of the human sera data, significance was determined by the nonparametric Mann-Whitney rank test. \**p*<0.05, \*\**p*<0.01, \*\*\**p*<0.001, \*\*\*\**p*<0.0001.

## Author Contributions

ZE and AA designed all experiments, interpreted the data, and wrote the manuscript. ZE and DO carried out all molecular biology, virus production and neutralization studies. KS and MF carried out animal studies and assisted with tissue analysis.

## Acknowledgements

This study was funded by NIH grants awarded to A.A. (R01HL089221, UG3AR07336, R01GM127708). We would like to thank StrideBio for providing the human serum samples for this study. We would also like to acknowledge Dana Elmore for her assistance with data analysis.

