## Supplemental Figures for "Rescuing AAV gene transfer from antibody neutralization with an IgG-degrading enzyme"

5148, MSRB3,

3 Genome Court,

Durham NC 27710, USA

**Supplementary Figures**

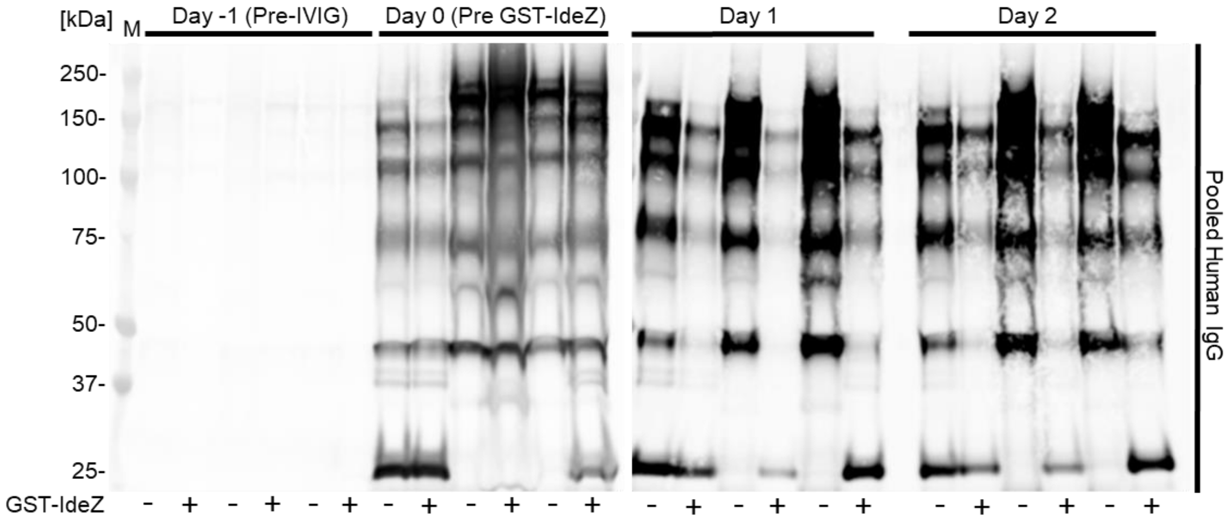


**Supplemental Fig 1. Kinetic analysis of IdeZ mediated cleavage of pooled human IgG *in vivo***. Mice were injected intraperitoneally with pooled human IgG. The same mice were injected intravenously 24 hours later with PBS (-) or recombinant GST-IdeZ (2.5 mg/kg) (+). Blood samples were taken 24 and 48 hours post injection and analyzed by SDS-PAGE under reducing conditions and probed with human IgG specific antibodies to analyze IgG cleavage.

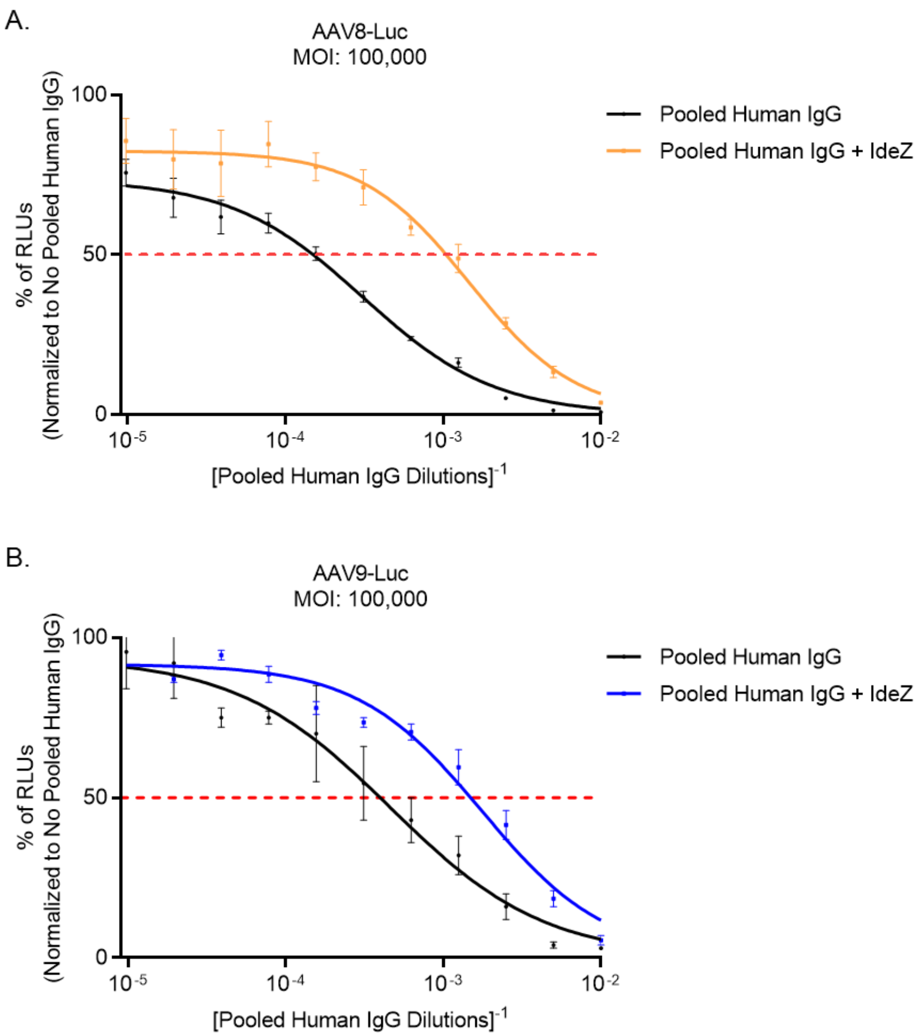


**Supplemental Fig 2. IdeZ mitigates human IgG mediated neutralization of AAV8 and AAV9 *in vitro*.**

Pooled human IgG was treated with PBS or GST-IdeZ (1ug). Samples were then serially diluted in twofold increments from 1:100 to 1:102,400 and then co-incubated with AAV8-Luc (**A**), or AAV9-Luc (**B**) *in vitro* (100,000 vg/cell) . The dotted red line represents NAb-mediated inhibition of AAV8/9 transduction by 50%. Solid lines represent relative transduction efficiencies of AAV8/9 incubated with IgG (black) and AAV8/9 incubated with IgG treated with GST-IdeZ (yellow,blue) at different dilutions. Error bars represent SEM (*n* = 2).

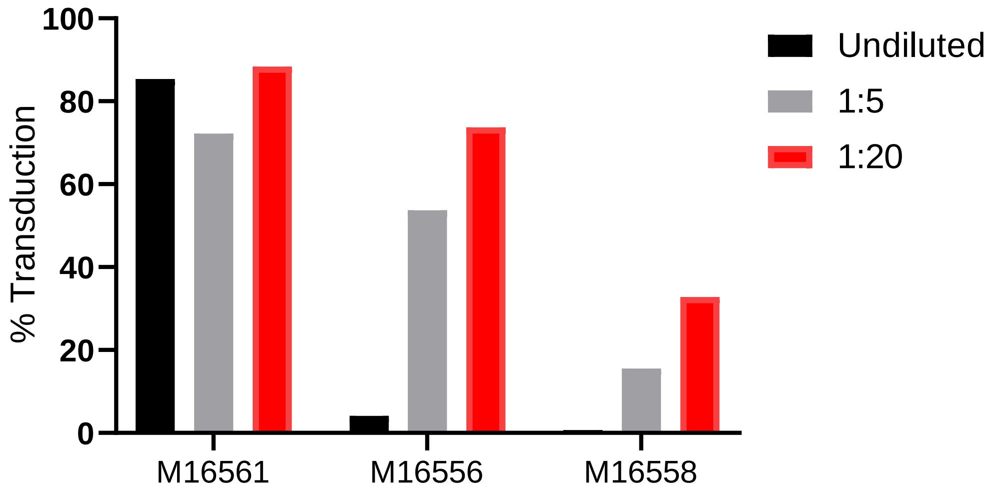


**Supplemental Fig 3. AAV9 neutralization profile of NHP sera.** NHP sera from three individual animals (M16561, M16556, M16558) were diluted from 1:5 to 1:20 and then coincubated with AAV9-Luc *in vitro* (100,000 vg/cell). Transduction levels were normalized to no serum control.

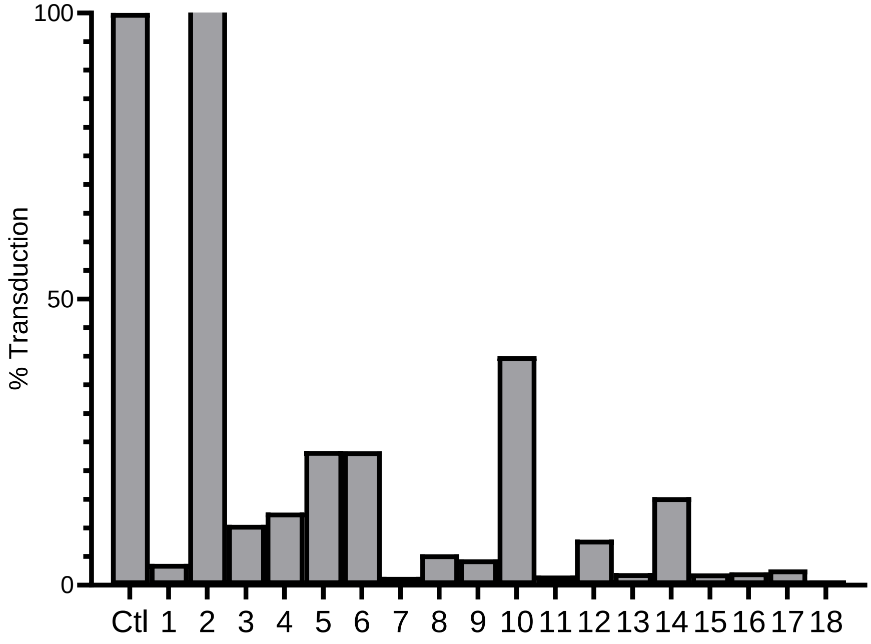


**Supplemental Fig 4. AAV9 neutralization profile of human sera.** Human sera from 18 individuals was diluted 1:5 and then coincubated with AAV9-Luc *in vitro* (100,000 vg/cell). Transduction levels were normalized to no serum control. Luciferase activity was determined at 24 hrs post-transduction. Detailed methods are described in Tse *et al.*, PNAS, 2017.

| Serum # | Lot # | Gender | Age | Race |
|---|---|---|---|---|
| **1** | BRH1499625 | Male | 37 | Hispanic |
| **2** | BRH1499636 | Male | 32 | Black |
| **3** | BRH1499659 | Female | 22 | Black |
| **4** | BRH1499617 | Male | 32 | Black |
| **5** | BRH1499665 | Female | 35 | Hispanic |
| **6** | BRH1499637 | Male | 37 | Caucasian |
| **7** | BRH1536076 | Male | 11 | Caucasian |
| **8** | BRH1499666 | Female | 27 | Black |
| **9** | BRH1499685 | Female | 35 | Caucasian |
| **10** | BRH1499676 | Female | 35 | Caucasian |
| **11** | BRH1499638 | Male | 30 | Caucasian |
| **12** | BRH1499620 | Male | 21 | Hispanic |
| **13** | BRH1536073 | Female | 16 | Black |
| **14** | BRH1499645 | Male | 34 | Caucasian |
| **15** | BRH1499679 | Female | 33 | Caucasian |
| **16** | BRH1499647 | Male | 35 | Caucasian |
| **17** | BRH1499669 | Female | 23 | Hispanic |
| **18** | BRH1499660 | Female | 35 | Hispanic |

**Supplemental Table 1. Demographic analysis of human serum used in this study.**

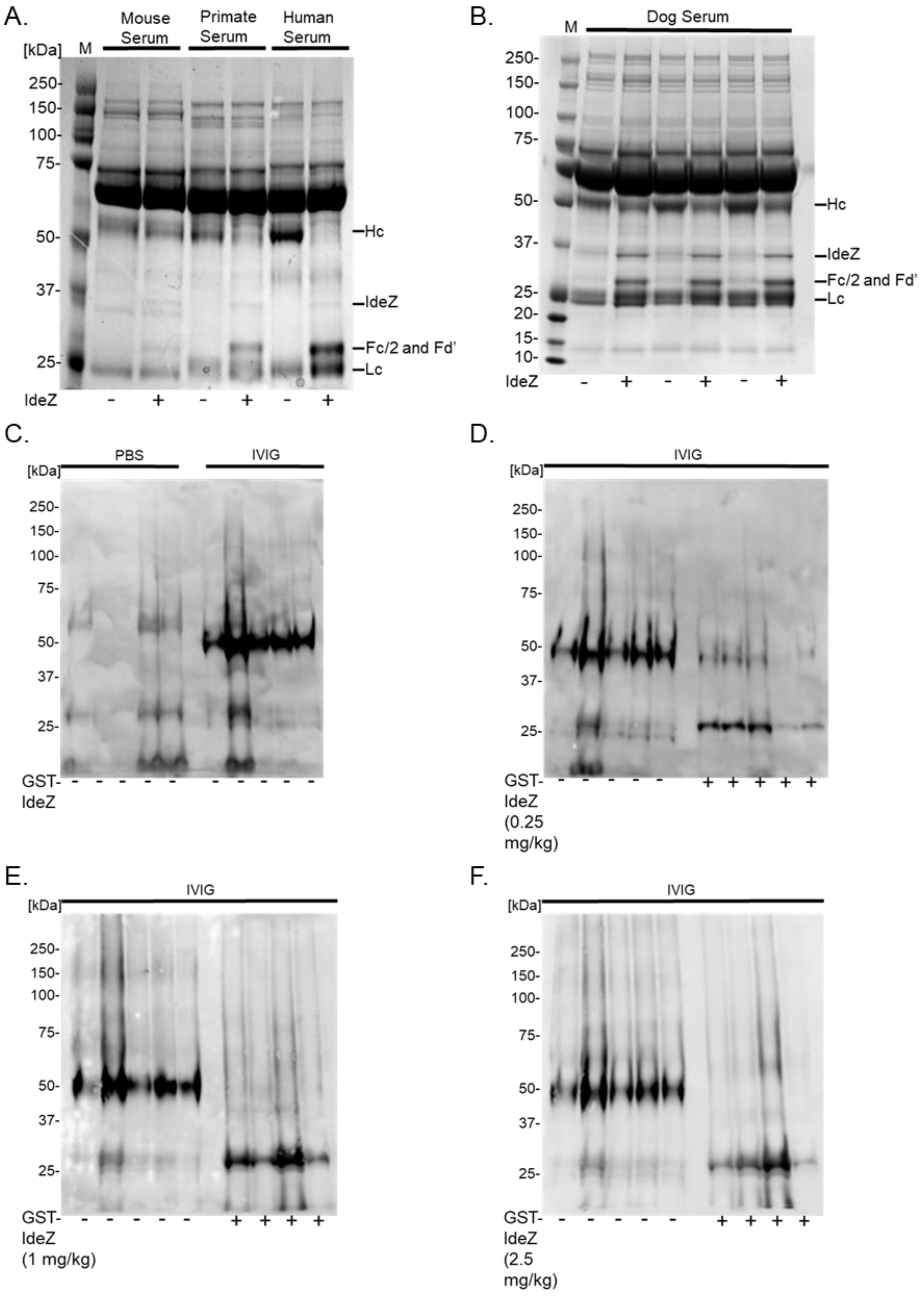


**Supplemental Fig 5. IdeZ biochemical analysis full size gels.**
